# Topological structure in human spatial representation revealed through drawing

**DOI:** 10.64898/2026.06.12.731853

**Authors:** Mahwish Kittur, Agnes Zhang, Nessa Bryce, Sami Yousif

## Abstract

Human spatial representations are often assumed to represent Euclidean properties such as length, distance, and angle. Here we test an alternative (but not mutually exclusive) possibility – that spatial memory is structured around topological relations. Across four experiments, adults and children memorized simple letter-like figures and reproduced them by drawing. Drawings showed systematic distortions of metric features, including strong biases of angles toward 90° and compression of line length towards an average value. In contrast, topologically critical features — such as T-junctions and holes — were reliably preserved, even relative to closely matched but topologically irrelevant features like L-junctions. These effects were magnified in a serial reproduction paradigm, in which participants iteratively generated new drawings from previous participant drawings: At the end of each mnemonic chain, figures converged on simplified topological structures as metric detail degraded. Similar patterns were observed in children aged five to eight years. Together, these findings suggest that topological relations may function as primitive building blocks of human spatial representation, with metric detail encoded inconsistently.

**Significance statement:** The iconic map of the London Underground is one of the most famous maps in history, yet something special about it goes unnoticed: it is not a veridical representation of space. Distances are arbitrary, and angles are presented only in coarse terms. Yet the ubiquity and appeal of such maps suggest that topological representation is intuitive — as if the mind is keen to receive information in exactly this way. Here, using drawing as a tool, we show directly that the most primitive form of spatial representation may be something akin to a topological skeleton. Remarkably, even children as young as five represent spatial structure in topological terms, with roughly the same fidelity as adults — pointing to an underappreciated building block of spatial representation.

---

Perhaps the foremost goal in cognitive science is to characterize the mental primitives underlying our rich representations of the external world. That is: We cannot possibly encode every detail about the external world into our memories, so the mind must selectively encode certain information in order to efficiently represent the features of the world that are most valuable. Thus, for every imaginable stimulus, a question arises about how it is encoded in the mind. What *content* is represented, and in what *format*? For instance: Is an image represented *pictorially* (see Kosslyn, 1994; Kosslyn et al., 1995) or *propositionally* (see Pylyshyn, 1973, 2002)? What primitive features are the building blocks of object representations – geons (Biederman, 1987), textons (Julesz, 1981, 1982), nonaccidental properties (see, e.g., Feldman, 1997, 2007; Kubilius et al., 2014; Kubilius et al., 2017), shape skeletons (Ayzenberg et al., 2019; Ayzenberg & Lourenco, 2019; Firestone & Scholl, 2014), or something else entirely? Is the cognitive map coordinate-based (Gallistel, 1990; O’Keefe & Nadel, 1978; Yousif & Keil, 2021) or graph-like (Chrastil & Warren, 2014; Warren et al., 2017; see also Yousif & Epstein, 2026)?

Here, we address a question of the same kind: We explore the basic units of human spatial representation that support our understanding of simple graphs and figures. Inspired by recent work on the subject (Yousif & Brannon, 2024, 2025; Yousif, Goldstein, & Brannon, 2025), we do so with a particular focus on *network topology* or *topological relations* – basic structures of graph representations like T-junctions, crosses, and holes (see Figure 1). Unlike prior work, however, we address this question by analyzing the features of simple spatial figures that are retained in simple drawings. The use of drawing as a tool allows participants to reveal directly the form of the representations in their mind. To understand how these mental representations develop throughout the lifespan, we analyze drawings from adults as well as children as young as five years old. To foreshadow: These drawings reveal a clear salience of topological features in participants of all ages, to an extent that is evident even from a cursory inspection of the drawings themselves.

**Figure 1.**
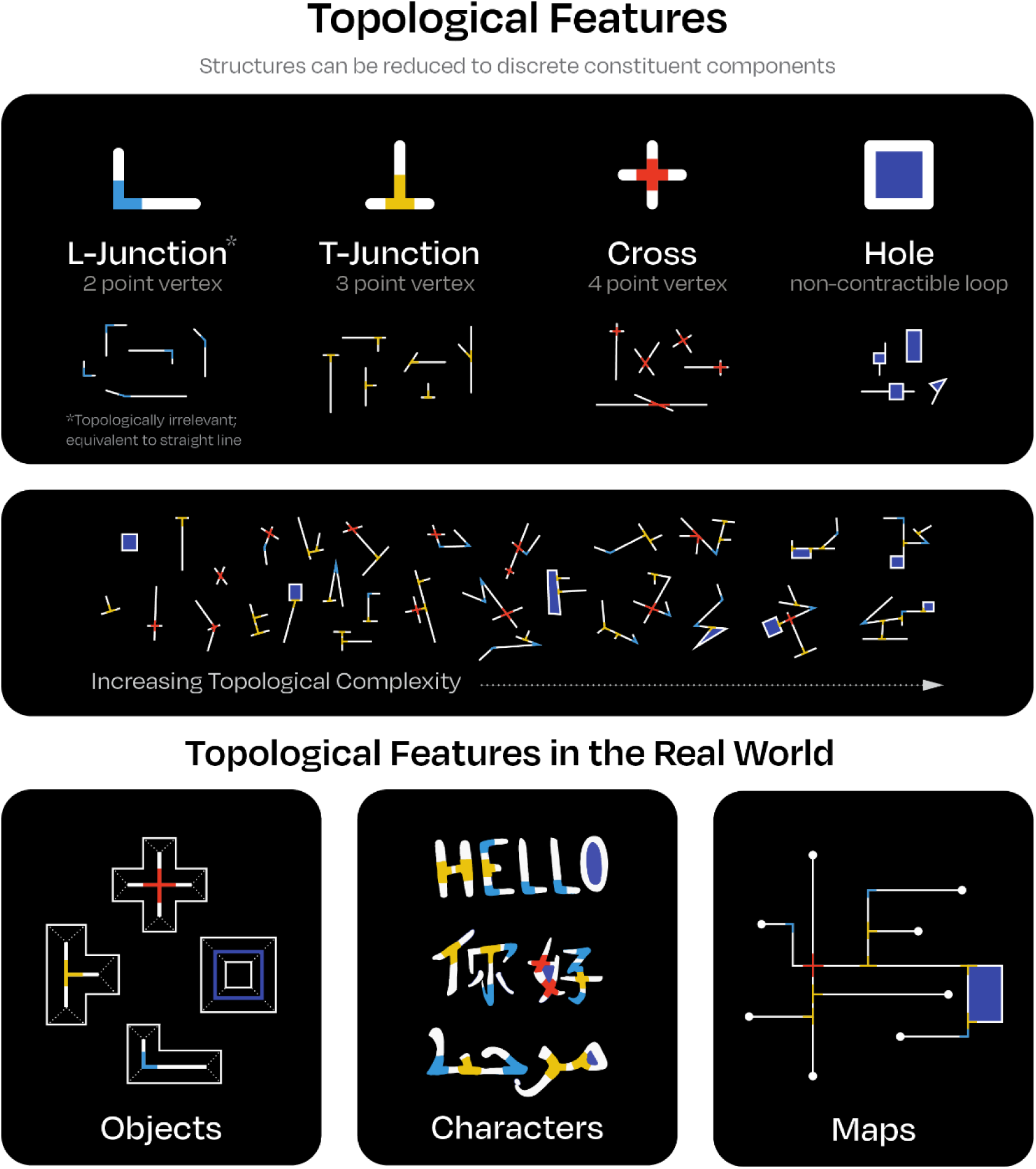
A visual depiction of various topological features like T-junctions, crosses, and holes. Note that L-junctions are a relevant contrast class but are not considered a topological feature. Forms with various topological structures are depicted. In the bottom panel, various examples of topological structure in ‘real world’ objects are shown.

## The building blocks of spatial representation

From the earliest stages of development into adulthood, most of what we’re taught about space reflects a particular *kind* of spatial representation. That is, we are overwhelmingly taught to think about space in Euclidean terms – with respect to lengths and angles, distances and directions. It is perhaps no surprise, then, that much work interested in spatial representation and spatial development has focused on Euclidean geometry (e.g., Dehaene et al., 2006; Lee, Sovrano, & Spelke, 2012; but see Huey et al., 2023; Kenderla et al., 2023).

For instance, Lee, Sovrano, & Spelke (2012) explored children’s use of Euclidean features in navigation, showing that children use some features (distance, direction) but not others (length, angle) to reorient following disorientation (but see Yousif & Lourenco, 2017). Ayzenberg & Lourenco (2020) refer to a “network for representing Euclidean geometry” (p. 1) that supports both navigation and object analysis. Each of these studies builds on the seminal insights of Hermer & Spelke (1994, 1996), who demonstrated that children use geometric features (e.g., distance, direction), but not non-geometric features (e.g., color, pattern), to reorient. Their work, in turn, drew from earlier work demonstrating that rats also exclusively use geometric information to reorient (Cheng, 1986), thus suggesting there is an evolutionarily ancient system of geometric representation that is shared by species across the animal kingdom.

In a similar vein, Dehaene and colleagues (2006) demonstrated that human adults and children without formal geometric education nevertheless possessed an intuitive understanding of geometric properties like parallelism, curvature, and distance – again supporting the suggestion that there may be “core” representations of geometry that are not merely products of formal education.

Yet Euclidean geometry is only one paradigm for thinking about space. Here, we focus on an entirely different paradigm: Topology. Whereas Euclidean geometry focuses on *precise* spatial information (i.e., exact distances, lengths, and angles) topology as a discipline is concerned instead with *coarse* spatial relations (i.e., the properties of spaces or objects that survive continuous deformations). The value and efficiency of topological representation is most evident via inspection of common transit maps, now popular across the globe (see Figure 1). What is notable about topological maps like these is that they intentionally distort the space. In reality, transit systems are not so perfectly designed as they seem in a topological map; the maps simply abstract away from all of the nitty-gritty Euclidean detail to focus on what matters most to passengers in these systems: The *relations* between the various lines and stations.

Another virtue of this form of topological representation is that it depends on a small number of discrete parts – features like T-junctions, crosses, and holes (see Figure 1) that can be combined to form networks of (virtually) infinite complexity. If you combine a hole and a T-junction in the right way, you may get a shape that resembles a “Q”; a hole with two T-junctions might form a shape resembling the letter “R”. These building blocks can be combined in a variety of other ways to generate essentially every character in every language on Earth. So, too, can they be combined to form simple graphical depictions of complex objects, maps, or concepts.

Prior work on human topological representation has focused largely on *object topology*, emphasizing, for instance, the distinction between rigid objects (i.e., rather than spatial networks) with and without holes (see, e.g., Chen, 1982). Differences in object topology influence visual perception (Chen, 1982; Chen, 1990; but see Rubin & Kanwisher, 1985), working memory (Wei et al., 2019), number perception (Franconeri et al., 2009; He et al., 2015), and object categorization (Kenderla et al., 2023), in human adults (Wei et al., 2019), human children (Chien et al., 2012; Kibbe & Leslie, 2016; Kenderla et al., 2023) and such distant species as bees (Chen et al., 2003). This is to say that topological structure is consequential – influencing many other psychological processes (see Figure 2).

**Figure 2.**
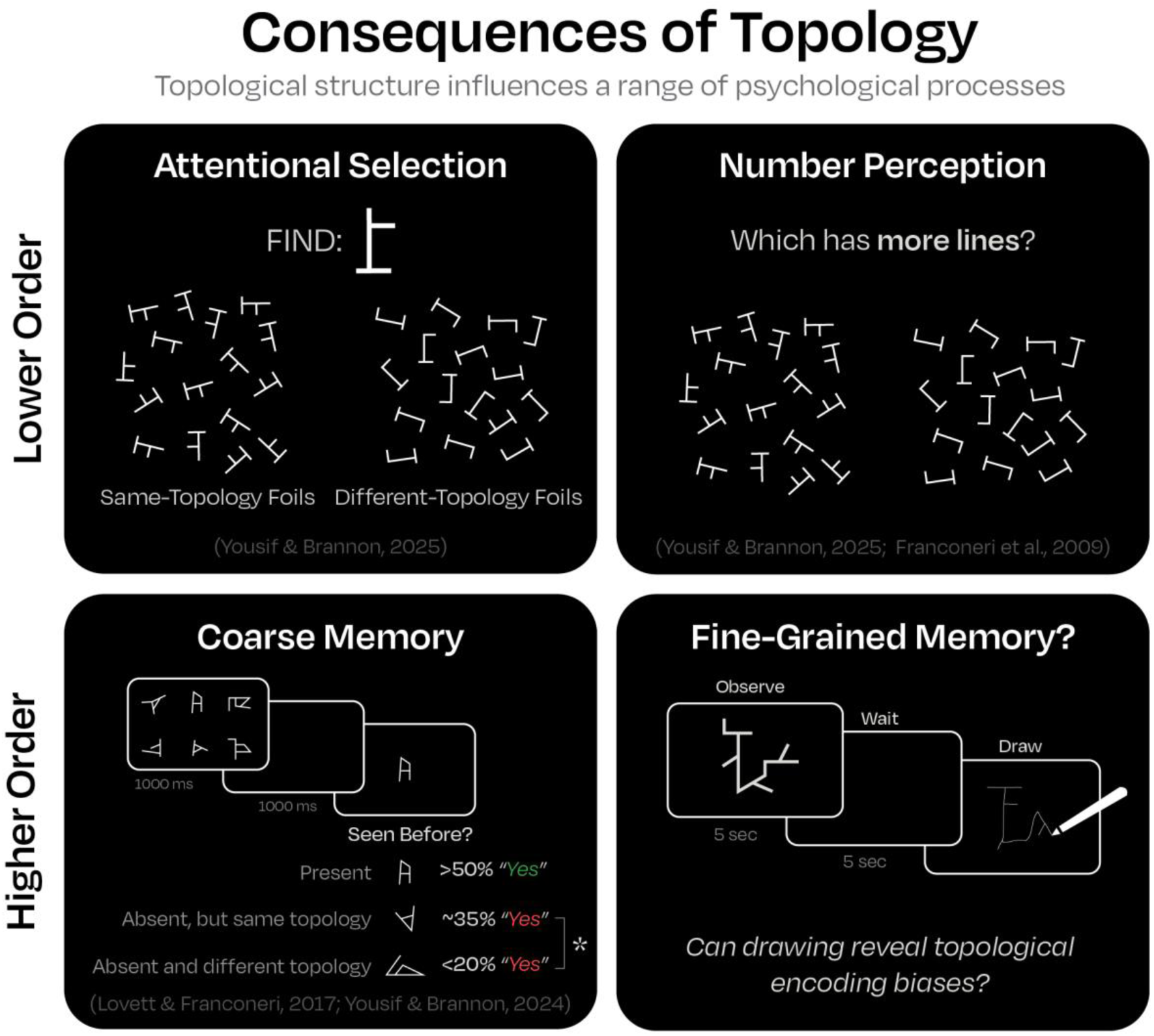
Consequences of topology. Topological structure influences various psychological processes from attention to number perception to memory.

In this study, we examine *network topology* (or *topological relations*, following the example of several recent papers; see, e.g., Shah & Yousif, 2026; Yousif, Goldstein, & Brannon, 2025; Yousif, Hu, & Brannon, 2026). Thus our analyses will focus on features of topological networks such as T-junctions, crosses, and holes (see Figure 1). Throughout this paper (as in prior work; see Yousif & Brannon, 2024, 2025), as a particularly stringent test of topological representation, we contrast T-junctions and L-junctions. In a certain light, both junction types are made up of two intersecting lines. Topologically speaking, however, T-junctions are functional whereas L-junctions are irrelevant. The latter reduces to a straight line. That is: If these junctions are imagined in segments of a maze, the T-junction presents the maze-goer with a choice, whereas the L-junction does not. From a topological perspective, it is the *choice* that matters; the choice reflects a meaningful, structural inflection point. On this view, any preferential representation of T-junctions over L-junctions would be consistent with a topological perspective. What is relevant is not the absolute rate at which either of these features is preserved, but the *relative* rate at which they are preserved.

To take a topological approach to spatial representation is not to imply that people *only* represent topological structure. It is undeniably the case that people represent non-topological information in at least some cases. Instead, the topological view proposes that topological information is *preferentially* represented as a relatively essential form of a spatial representation and that precise, metric information (as well as other non-topological information) is perhaps superimposed on top of a coarse, topological representation. In this way, the topological view predicts that in memory, non-topological information may be systematically distorted or lost, while topological information is robustly preserved.

There are many hints of topological representation in prior work. It is known, for instance, that participants experiencing a new spatial environment draw out maps of that environment with significant angular distortions. Namely, people are predisposed to draw all angles as closer to 90° than they really were (Byrne, 1979; Moar & Bower, 1983). This makes sense if all intersections are being represented as nodes in a network; in a topological representation, only discrete intersections matter, not their precise angle (see also Chrastil & Warren, 2014; Warren et al., 2017). Similar biases have been observed in non-navigation contexts, both in adults and children (Bremner & Taylor, 1982; Ibbotson & Bryant, 1976), though there is some debate about whether/when children first exhibit a 90-degree bias (cf., Ibbotson & Bryant, 1976; Izard & Spelke, 2009). Numerous other studies have demonstrated perceptual prioritization of the cardinal axes compared to the oblique axes, perhaps also reflecting a sort of perpendicularity preference (see, e.g., Rademaker et al., 2017; Yousif, Chen, & Scholl, 2020; Yousif & McDougle, 2024). Relatedly, prior work has shown robust distortions of length/distance (because of differences in network structure) when participants are asked to draw maps and other map-like stimuli (Klippel et al., 2005; McNamara et al., 1984). Some work has even shown that the presence of topological features like T-junctions in drawings is significantly related to the memorability of that image (see, e.g., Han, Rezanejad, and Walther, 2023). All told, there is a considerable body of work from both the spatial cognition literature and the broader vision science literature which could be viewed as consistent with a topological view of spatial cognition.

## Drawing as a “tool” in cognitive science

The aim of this paper is to analyze topological representation and Euclidean representation in tandem by accessing the form of human spatial representations directly – via drawing.

Drawing is a critical part of human history, with the first known drawings dating back hundreds of thousands of years (Joordens et al., 2015), if not longer (Le Tensorer, 2006). Indeed, these drawings offer considerable insight into the foundations of human cognition: The universality of drawing across culture, age, and time suggests an innate capacity for representing (certain kinds of) complex information in a visual form (see, e.g., Sablé-Meyer et al., 2022). So too does drawing have a rich history within psychology and cognitive science. Nearly one hundred years ago, Goodenough (1926) created the “draw-a-person” test which supposedly assessed both personality and intelligence. Bartlett (1932) showed how memories become distorted through memory and transmission via drawing, using the now-storied method of serial reproduction. Piaget used drawings to provide insight into the child’s mind in many ways throughout his career, but most notably in his work with Inhelder exploring children’s spatial representation (Piaget & Inhelder, 1948). In fact, partly on the basis of such work, Piaget and Inhelder proposed that children’s spatial representations were fundamentally topological in nature.

In more recent years, drawing has been used as a tool in the study of spatial neglect (Agrell & Dehlin, 1998), cognitive and memory disorders (Brodaty & Moore, 1997; Gainotti et al., 1983), imagery (Bainbridge et al., 2021), visual perception (Fan et al., 2018), visual memory (Cardenas-Miller et al., 2025; Fernandes et al., 2018; Hall & Geng, 2023; Wammes et al., 2016), memorability (Han et al., 2025), and cognitive development (Dillon, 2021, Long et al., 2024). In all these cases, the value of drawings is obvious: Rather than asking participants pointed questions, they can be given a (literal) blank canvas upon which to reveal the form of their mental representations. Drawings are a tool for revealing, in perhaps the most direct manner possible, the structure of a representation (see also Bainbridge et al., 2025; Fan et al., 2023).

## Current study

Here, in four experiments, we use drawing as a tool to understand topological representation in adults and children. Experiments 1 and 2 were both focused on adults; we used two different stimulus sets specifically designed to assess certain spatial features. Experiment 3 took inspiration from Bartlett’s (1932) classic memory studies: Groups of participants ‘serially reproduced’ drawings one after the other, resulting in chains of figures that were gradually distorted. Experiment 4 focused on children’s drawings, using the same analytical approach as the adult experiments. In all of these cases, participants were simply shown a figure for several seconds and then asked to draw it from memory a few seconds later (see Figure 3).

**Figure 3.**
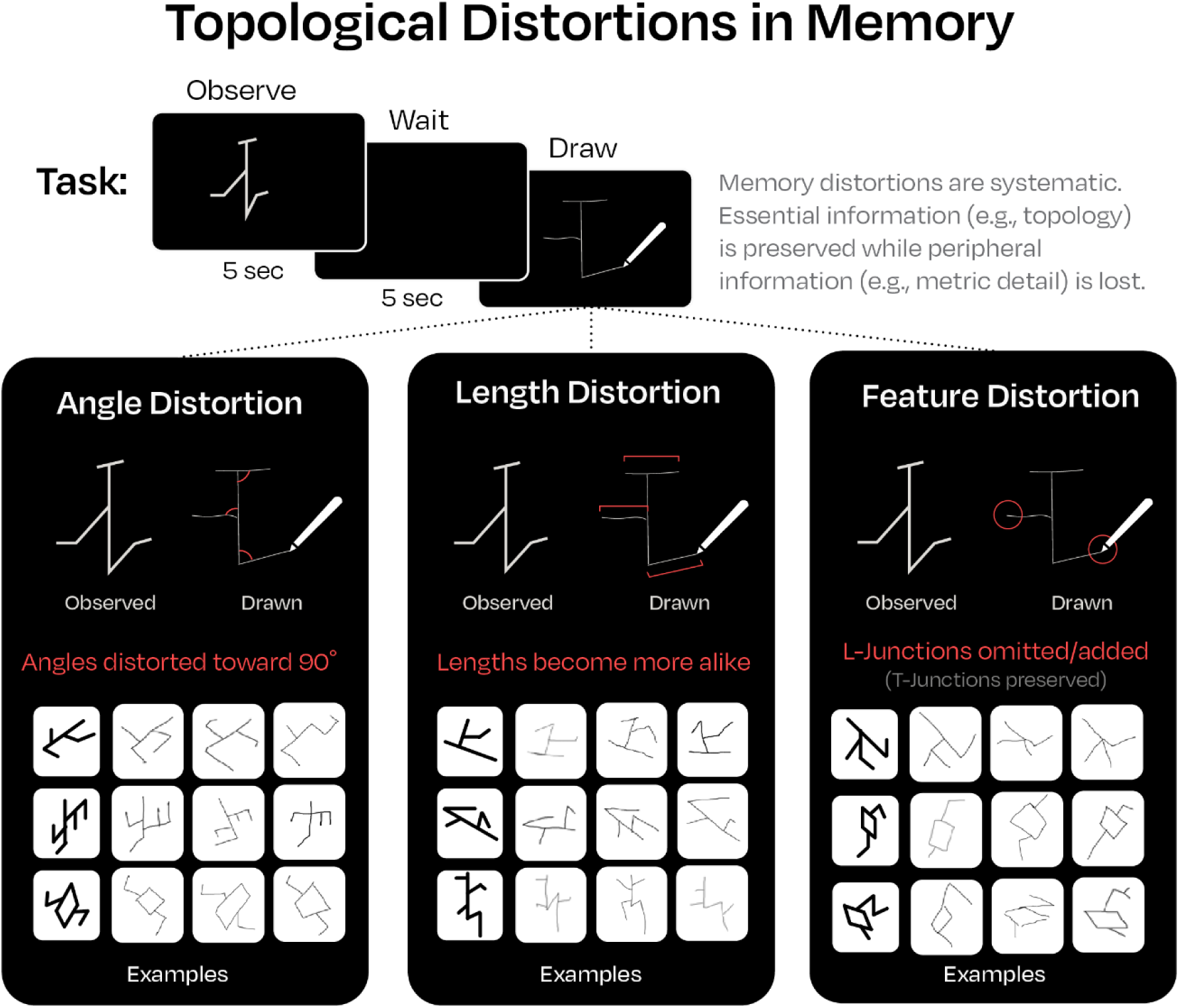
A simple depiction of the drawing paradigm, as well as depictions of the three key distortions measured in this paper. Representative examples of each distortion, drawn from real adult drawings, are shown.

Across these studies, we focus our analyses on a few key aspects of the drawings (see Figure 3). First, we are interested in the relative preservation of distinct topological features like T-junctions and holes. Most importantly, we contrast T-junctions and L-junctions as a particularly strict test of the topological view (see above). Second, we are interested in the relative distortion of, or loss of information about, metric properties like length and angle. Specifically, we are interested in whether length and angle information is systematically distorted to a ‘default’ value (e.g., 90 degrees, an average line length). Note that in both of these cases, these analyses are strictly hypothesis driven. The predictions that T-junctions will be preferentially preserved and that length and angle information will be distorted are key predictions of the topological view; thus, the observation of such patterns should be seen as strong validation of it.

## Results

### Experiment 1: Angular distortions

In a first experiment, adult participants were presented with a simple memory task. They were shown letter-like stimuli for five seconds each, asked to memorize them, and then, after a short delay, draw them from memory to the best of their ability (see Figure 3). We tested two hypotheses. First, we investigated the distortion of angular information. Since precise angles are topologically irrelevant, we asked whether participants would be biased towards representing all angles as closer to 90°. Second, we tested whether participants were more likely to distort T-junctions or L-junctions. Since L-junctions are not topologically relevant (as they have the same topological structure as a line), we asked whether participants would be more likely to make errors for L-junctions than T-junctions. For more information, see *Methods*.

Results can be seen in Figure 4 (A, B, & D). First, we measured the drawings for evidence of angular biases. We measured both the proportion of drawings with angles biased towards 90° (i.e., the proportion of all measured angles that had a lesser absolute distance from 90 degrees compared to the original angle; see *Methods*), as well as the actual average bias. On average, each participant drew 72.3% (SD=7.4%) of the measured angles as closer to 90° (*t*(49)=21.38, *p*<.001, *d*=3.02; see Figure 4A), by an average amount of 12.11° (SD=3.21°; *t*(49)=26.71, *p*<.001, *d*=3.78; see Figure 4B). Indeed, inspection of Figure 4B reveals that all angles, regardless of their original magnitude, are overwhelmingly biased towards 90°. Overall, this same pattern of bias towards right angles was observed for nine of the ten unique stimuli tested. Note that these findings are consistent with prior work showing that both for 2D and 3D spatial representations, angles are robustly distorted toward 90° (Bremner & Taylor, 1982; Ibbotson & Bryant, 1976). However, prior findings have seen such patterns as evidence of some sort of 90° bias, whereas we view these patterns as evidence of a topological representation of space that does not encode precise angular information in the first place.

**Figure 4.**
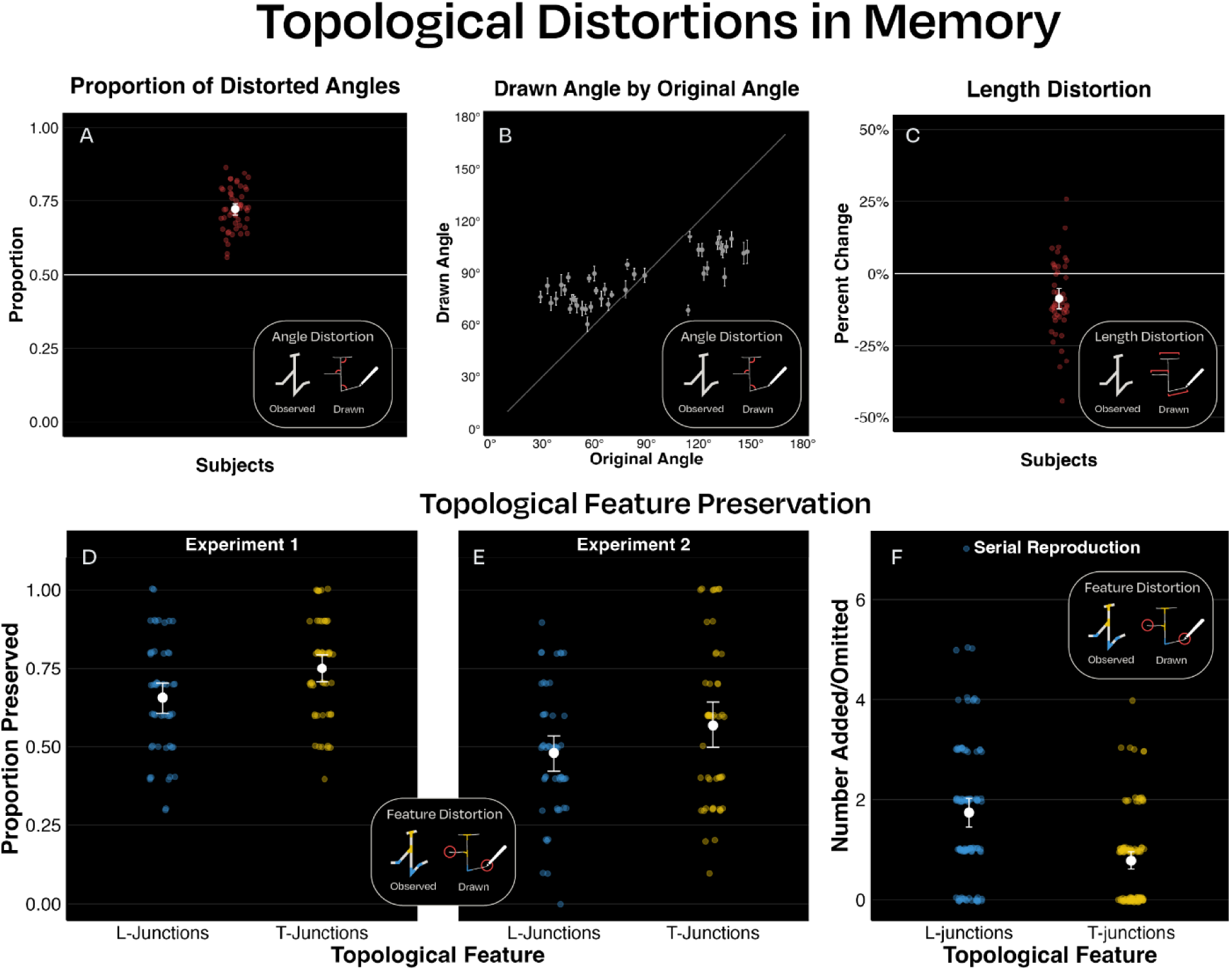
Results from Experiments 1-3. (A) The proportion of angles biased towards 90 degrees for each participant. (B) The reproduced angle as a function of original angle, for each of the 42 unique angle sizes that were measured across the 10 objects. (C) The average distortion of the relative line lengths for each participant. Negative values indicate that the lines were drawn as closer in length. (D) Relative preservation of L-junctions versus T-junctions in Experiment 1. (E) Relative preservation of L-junctions versus T-junctions in Experiment 2. (F) Relative preservation of L-junctions versus T-junctions in Experiment 3. Note that in this final case, we plot the number of added/omitted features *across the entire sequence*. Thus, the axis in this panel is effectively flipped relative to the previous two panels.

Further evidence of topological representation comes from analysis of the topological features themselves. As a strong test of the topological view, we focused on the distinction between L-junctions and T-junctions. L-junctions and T-junctions, while visually similar, are topologically distinct. Critically, T-junctions are a meaningful topological feature, whereas L-junctions, on a strict interpretation, are not topologically relevant (in the sense that they are reducible to a straight line, via bending). Thus, we reasoned that, if participants represent something akin to the true topological structure, they may preferentially preserve T-junctions rather than L-junctions in their drawings. Thus, we compared the relative likelihood of preserving the correct number of each feature. As a topological perspective predicts, participants were indeed more likely to recreate the correct number of T-junctions (M=75%, SD=16.2%) than L-junctions (M=65%, SD=18.4%), *t*(49)=5.11, *p*<.001, *d*=.72. Examples of these types of distortions can be seen in Figure 3. Together, these results indicate that (A) topological structure does seem to influence what features are and are not retained in drawings, and (B) topologically irrelevant information, like angular information, is consistently lost.

The analysis conducted here comparing T-junctions and L-junctions is meant to be an especially strict test of the topological view. One thing not captured by this analysis, however, but obvious from inspection of the raw drawings themselves, is that topological features across the board are overwhelmingly preserved. Holes, for instance, virtually never disappear from drawings (<1% of the time).

But do these patterns reflect errors of memory, or poor drawing ability? While it is unclear how poor drawing would explain a *bias* in angular memory or the tendency to forget certain kinds of junctions, we ran a supplemental reproduction task in which participants were asked to draw while viewing the image side-by-side. Putatively, this design isolates errors introduced from drawing ability alone. These data are included as Experiment S1 in the raw data on our OSF page and are described in more detail in the supplemental material there. To summarize those data succinctly: There was no angular bias, virtually no topological errors, but interestingly, a similar degree of distortion for line lengths (see Experiment 2). Even cursory examination of the drawings themselves (also visible on our OSF page) reveals striking differences between the drawings reproduced live and the ones reproduced from memory.

### Experiment 2: Length distortions

The stimuli used in Experiment 1 were specifically designed to test for angular distortions. However, angles are not the only Euclidean feature we might expect to be distorted. If people are truly representing these objects with respect to their topological structure first and foremost, we might also expect that, e.g., the lengths of lines are similarly distorted (e.g., such that all line lengths become ‘pulled’ toward a central value). Thus, our second experiment had two aims: First, we wanted to replicate the T-junction vs. L-junction effect from the first experiment. Second, we wanted to analyze distortions of line lengths. To accomplish both of these goals, we designed a new set of stimuli specifically optimized for addressing these questions.

Results can be seen in Figure 4 (C & E). Whereas the previous stimuli were designed to test angular biases, these stimuli were specifically designed to test whether line lengths are distorted towards an average value. Our analyses focused on the line segments in each drawing that were created to have a precise 4:2:1 ratio. For each drawing, we measured the relative length of each of those lines, computing the ratio between the longest line and the middle line, as well as the middle line and the shortest line. To get a final value for each stimulus, we calculated the geometric mean of those two ratios. In practice, this means that ratio values below 2.0 are indicative of compression towards an average value, whereas ratios above 2.0 would be indicative of expansion.

Overall, the (geometric) mean of the drawn line length ratio for each participant was 1.86 (SD=.21), significantly below the original ratio of 2.0, equivalent to an 8.6% reduction on average (computed as (r − 2)/r for ratios below 2 and (r − 2)/2 for ratios above 2; see Figure 4C), *t*(49)=4.87, *p*<.001, *d*=.69. These data indicate that, like angle information, length information is distorted towards a ‘prototypical’ value.

In addition, we once again analyzed whether T-junctions were preserved at a higher rate than L-junctions. We found that the former were maintained 56.8% (SD=25.6%) of the time whereas the latter were maintained only 48.8% (SD=23.2%) of the time, *t*(49)=2.72, *p*=.009, *d*=.38. Collectively, these data provide converging support for the topological perspective.

### Experiment 3: Serial Reproduction

Both of the prior experiments reveal biases consistent with topological representation. Even so, these biases may be considered relatively subtle. What if we could magnify these biases so that they were apparent not only from careful data analyses, but via mere inspection of the drawings themselves? In this experiment, we did so by employing a serial reproduction paradigm (Bartlett, 1932). In this design, a first participant was shown a drawing much like had been done in the prior experiments. Then the drawing that they output would be used as the *input* for a second participant, whose output would in turn be used as the input for a third participant, and so on, for ten iterations (see Figure 5). The idea behind this approach is that the cumulative errors across a chain will cause the figures to slowly approach the most primitive form of the representation.

**Figure 5.**
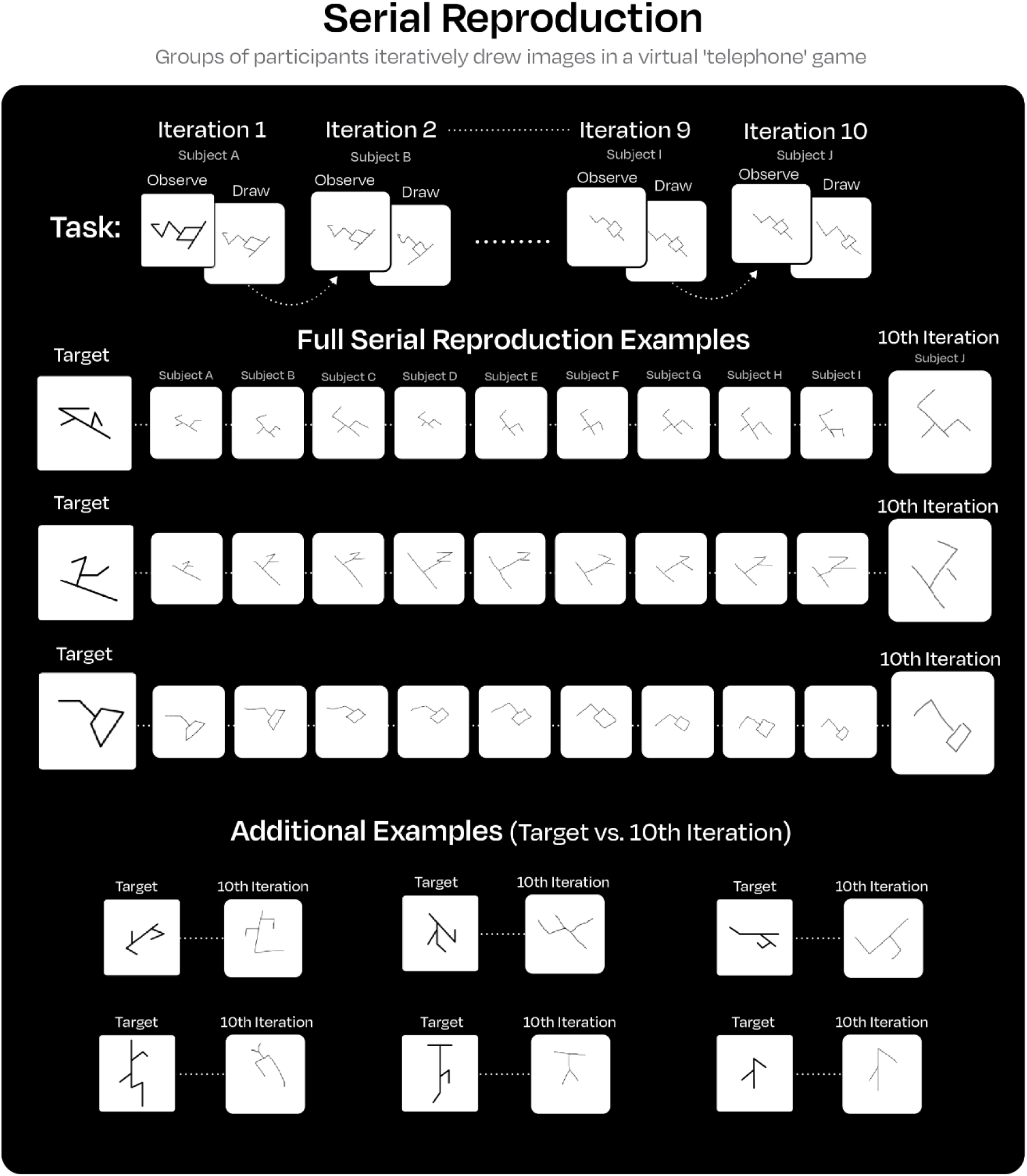
A depiction of the serial reproduction paradigm, as well as examples of partial and complete ‘chains’ of drawings. As is evident from these drawings: Length and angle information were robustly distorted throughout the chain, while topological structure was largely preserved. Note that each one of the ten unique stimuli is depicted exactly once.

Results can be seen in Figure 4F and Figure 5. As with the adult data in Experiment 1, we first analyzed the drawings for evidence of angular biases. On average, the final drawing in each chain had angles drawn nearer to 90° by an average amount of 17.07° (SD=16.55°; *t*(99)=10.32, *p*<.001, *d*=1.03), slightly more than the bias observed in Experiment 1.

We also examined whether lengths were biased towards an average value, as in Experiment 2 (but only on the subset of five stimuli specifically designed with 4:2:1 length ratios). Three of the fifty chains were excluded from the analysis because the coders could not agree on measurements. Overall, the (geometric) mean of the final line length ratio for each chain was 1.38 (SD=.35), equivalent to a 50% reduction, *t*(46)=8.11, *p*<.001, *d*=1.18. Note that the length biases are significantly larger than those observed in Experiment 2, likely because these biases were able to accumulate and grow larger over the course of the 10 reproductions.

Finally, we examined whether the final drawings in each chain were more likely to have retained the original number of T-junctions as opposed to L-junctions. We found that T-junctions were marginally more likely to be preserved by the end of the sequence. Of the 100 sequences, T-junctions were preserved 46% of the time, whereas L-junctions were preserved only 36% of the time. On average, .93 L-junctions were added or removed in the final drawings whereas only .74 T-junctions were, *t*(99)=1.79, *p*=.076, *d*=.18.

As an even stronger test of this account, we also analyzed changes in the number of topological features *throughout* each sequence. This analysis is critical because (as earlier results revealed), L-junctions are both more likely to be added and more likely to be removed from drawings than T-junctions. Thus, analyzing only the final drawings leaves out meaningful changes that may have occurred throughout each sequence. On average, participants added or removed L-junctions 1.74 times (SD=1.41) compared with only 0.78 times for T-junctions (SD=.92), *t*(99)=6.93, *p*<.001, *d*=.69 (see Figure 4F). Additionally, they added or removed only .22 crosses (SD=.58), significantly less than the T-junctions, *t*(99)=6.10, *p*<.001, *d*=.61.

Empirically, these results reinforce the findings of Experiments 1 and 2 and further validate the topological perspective. Visually, these results reveal even more striking topological biases. Even cursory inspection of the chains of drawings (see Figure 5) reveals many of the patterns reported here: Angles often quickly converge on 90°, and line lengths gradually gravitate towards one another. Despite these biases, it is also plain to see that the overall topological structure is often well-preserved, sometimes even despite significant distortions of the other spatial features.

### Experiment 4: Angular Distortions with Children

Decades ago, Piaget and Inhelder (1948) proposed that children’s spatial representations are fundamentally topological. So far, we have shown clear signs of topological representation in adults’ drawings. If Piaget and Inhelder were right, we might expect that children exhibit even clearer evidence of topological structure in their drawings (and, relatedly, clearer evidence of distortions of geometric features). Here, we use the same approach as Experiment 1, except that we tested five-to eight-year-old children.

Results can be seen in Figure 6. As with the adult data in Experiment 1, we first analyzed the drawings for evidence of angular biases. On average, each participant drew 77.1% (SD=8.2%) of the measured angles as closer to 90° (*t*(125)=37.31, *p*<.001, *d*=3.32), by an average amount of 14.01° (SD=4.49°; *t*(125)=35.05, *p*<.001, *d*=3.12). Age was not a predictor of either of these effects (magnitude: *b*=0.39, SE=0.41, *t*(124)=0.94, *p*=.35; proportion: *b*=.003, SE=0.008, *t*(124)=0.38, *p*=.70).

**Figure 6.**
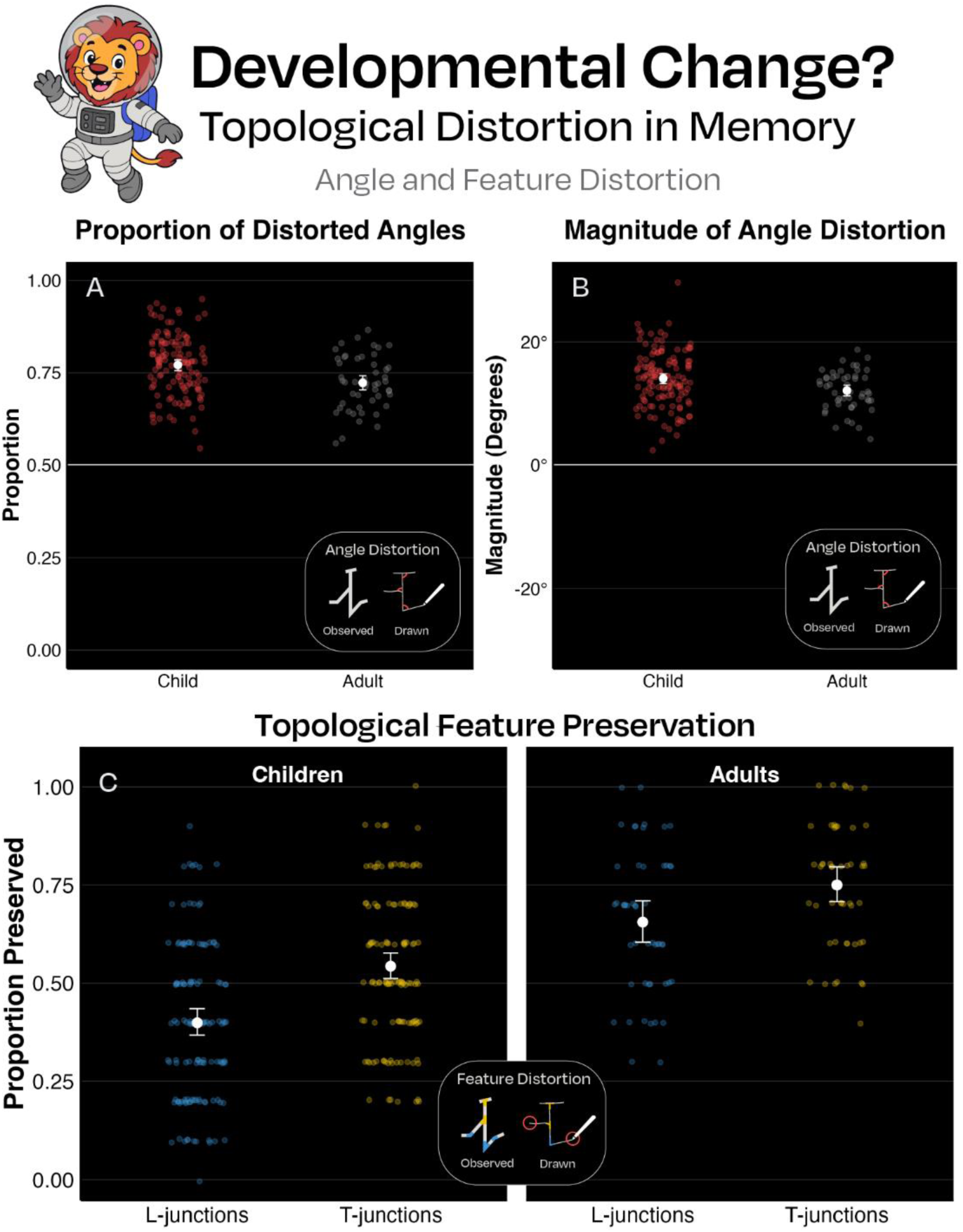
Leo the lion and the results from Experiment 4. (A) The proportion of angles biased towards 90 degrees for each participant, broken down by age. Adult data from Experiment 1 are plotted as a comparison. (B) The average magnitude of angular bias in each participant, in degrees, broken down by age. Positive values indicate bias in the direction of 90 degrees (see Methods). Adult data from Experiment 1 are plotted as a comparison. (C) Relative preservation of L-junctions versus T-junctions. Adult data from Experiment 1 are plotted as a comparison.

Like adults, children were also more likely to draw the correct number of T-junctions (M=54.3%, SD=19.2%) than L-junctions (M=39.8%, SD=19.8%), *t*(125)=8.59, *p*<.001, *d*=.77. Age predicted the tendency to preserve both L-junctions (*b*=0.094, SE=0.016, *t*(124)=5.79, *p*<.001) and T-junctions (*b*=0.10, SE=0.015, *t*(124)=6.57, *p*<.001) but not the difference between the two (*b*=0.006, SE=0.017, *t*(124)=.36, *p*=.72). This is consistent with the fact that the overall rates of feature preservation were higher in the adult sample.

While children’s drawings are improving throughout early childhood, their topological biases remain consistent from the earliest ages tested here. These patterns coincide with prior work showing striking stability in simple object concepts from four years old into adulthood (see Yousif et al., 2025). Thus, these findings suggest that whatever primitive representations or processes guide spatial representation are early developing, present from approximately the age of four onward. A question arises, then, about whether even young children represent simple spatial structures in similar ways, and, possibly, whether there are spatial primitives that are innate. Future work may address this possibility by probing sensitivity to topological structures in infancy.

We view the preservation of topological features like holes and T-junctions as consistent with the distortion of Euclidean features like length and angle (in the sense that coarse topological information is selectively preserved in memory while the precise metric information is lost). However, even if one rejects this premise, it is nevertheless true that children show the same fundamental pattern of memory errors as adults, thus raising questions about what underlying representational system is responsible for these stable patterns of responses.

What role might motor abilities play in these findings? We have no independent measure of fine motor skill, but we assume it continues developing across the tested age range. It seems unlikely, however, that differences in motor ability explain the observed results, for two reasons. First, age did not predict any of the key effects. Second, and perhaps more importantly, reduced motor ability should introduce *noise* into drawn figures, not *bias*; there is no clear reason it would bias reproductions toward a particular point in representational space (e.g., toward 90-degree angles). While we acknowledge that motor abilities may play some role in tasks like these, we see little reason to think the core conclusions of this experiment could be explained by such differences.

## General Discussion

Memory is fallible. The visual system receives rich information, but only some of that information will be retained with any fidelity. Yet those parts that do remain are telling: They reveal the most primitive information that we encode. Here, we have shown that, when remembering simple spatial structures, people seem to encode not precise lengths or angles, but instead coarse *topological relations*. Indeed, despite robust distortions of both angle (Experiment 1) and length (Experiment 2), topological features like T-junctions were relatively well preserved (even compared to topologically irrelevant junctions, like L-junctions; see Experiments 1 and 2). Drawings from a serial reproduction paradigm made each of these patterns even more clear: By the end of each chain of drawings, line lengths, and especially angles, were overwhelmingly distorted, while the overall structure of the original shapes was surprisingly well preserved (Experiment 3). Similar distortions were observed in children as young as five years old (Experiment 4), suggesting that these biases reflect early developing spatial knowledge. That said, we did not find any convincing evidence that children were more topologically inclined than adults. Whereas Piaget and Inhelder (1948) described an intuitive tendency for topological representation that erodes throughout development, the evidence here is more consistent with a stable developmental trend (see also Yousif et al., 2025). Collectively, these data suggest that basic topological structures may serve as primitive building blocks of visuospatial representation early in life and throughout the lifespan.

Whereas prior work on topological representation has relied on classic experimental designs, here we employed a maximally open-ended approach. Rather than constraining the choices participants could make, we allowed them to show us directly what they represented in their minds. This approach has several advantages. Chief among them is the fact that we can then study all aspects of the representation in tandem. Within a single drawing, we can see both the preservation of topological structure and the distortion of metric detail, as well as, conceivably, any other systematic bias of representation. Further, this approach allows us to see how these distortions evolve and grow as the figures are communicated from person to person (via serial reproduction; see Experiment 3). Because the drawing task requires no intervention from the experimenter whatsoever (either at the design or test stages), these data paint a particularly clear picture of the nature of human spatial representation.

That said, it is worth separating two distinct patterns in our results. The first pattern concerns the *distortion* of angle and distance. We believe the present data provide compelling evidence that metric features are represented with poor fidelity such that they degrade in memory over time. The second pattern concerns the *preservation* of topological features like T-junctions and holes. This pattern may be broadly interpreted in one of two ways. A weak interpretation of this pattern is that topological structure just happens to be preserved as metric information is degraded — but not because the mind encodes topological structure *per se*. On this interpretation, while the present data may provide strong evidence that certain information (e.g., metric information) is encoded noisily, they may provide only a hint about the true structure of the representation. A stronger interpretation is that people encode a *bona fide* topological representation such that the mind explicitly represents different junctions/parts and the relations between them. The drawing data, on their own, do not enable us to distinguish between these two interpretations. In what follows, we advocate for a slightly stronger interpretation, but only in light of (i) converging evidence exploring these exact kinds of topological representations (see, e.g., Yousif & Brannon, 2024, 2025; Yousif, Hu, & Brannon, 2026), and (ii) additional evidence of similar representational distortions in very different tasks (e.g., in drawing tasks following navigation; see Moar & Bower, 1983; Byrne, 1979).

### Topological structure as a primitive representation

Children begin learning about Euclidean geometry from an early age. By the end of elementary school, many students will have learned about geometric concepts like perimeter, area, and volume. Long before that — before they even enter the classroom — they learn informally to distinguish shapes based on the sizes of their angles and the numbers of their sides. Even earlier, still, they are said to possess a natural affinity for certain Euclidean geometric concepts: Newborn infants notice changes in shape (Slater et al., 1991), suggesting that humans may have some “core” geometric intuitions which then serve as a foundation for the acquisition of geometric knowledge throughout the lifespan (Dehaene et al., 2006). Indeed, geometric knowledge is predictive of achievement in mathematics, science, engineering, and more (Cheng & Mix, 2014; Clements et al., 2022; Newcombe, 2010; Vallortigara, 2012; Verdine et al., 2017). Perhaps unsurprisingly, then, much work investigating primitive spatial representations has taken a Euclidean approach (see Dehaene et al., 2006; Dillon, Huang, & Spelke, 2013; Izard, Pica, & Spelke, 2022; Izard & Spelke, 2009; Sinclair et al., 2016; Yousif & Lourenco, 2017).

However, there are abundant reasons to think that topological representations are a core part of human spatial representation. For one thing, there is considerable work showing that principles of object topology influence visual perception (Chen, 1982, 1985, 2005), object understanding (Kenderla et al., 2023; Kibbe & Leslie, 2016; Yousif & Brannon, 2024), and even numerical perception (Franconeri et al., 2009; He et al., 2015; Yousif & Brannon, 2025). Such work spans all sorts of populations from human adults, to human children and infants (Chien et al., 2012; Kenderla et al., 2023; Kibbe & Leslie, 2016; Yousif et al., 2025), to bees (Chen et al., 2003). More concretely, topological representations are a common part of everyday life for hundreds of millions of people across the world. Topological maps (see Figure 1), common to transit systems in almost every major city in the world, are valued by passengers precisely *because* they abstract away from metric detail to instead represent topological structure. The predisposition in favor of topological maps might be viewed as a critical clue that this information is presented in a manner that our minds are keen to receive it — perhaps because this is an essential default form of spatial representations.

There are also several independent reasons to think that the building of large-scale cognitive maps may depend on topological representation. First, there is the simple fact that, whether we realize it or not, we predominantly navigate the world through graph-like structures — city streets, subway systems, a web of interconnected airports. Our experience with the spatial world is far removed from the open world itself; we experience it, and traverse it, in highly predictable, structured ways. It would surely benefit us, then, to be able to represent those large-scale spaces in a comparable form. Second, classic work in spatial cognition has shown biases in map-drawing of navigable spaces similar to the biases observed here. Individuals make the same systematic distortions in angle and length/distance in their reproductions of space as they do in their reproductions of form. For example, people consistently estimate angle measures as closer to 90° in their mental representations of spaces, even when true angle measurements differ greatly from 90° (Moar & Bower, 1983; Byrne, 1979). Similarly, reproductions of distance reveal distortions in relation to various other factors, such as the number of turns, landmarks, and other heuristic information (Roberts et al., 2011; Bugmann & Coventry, 2008; Costa & Bonetti, 2018). These common distortions indicate a possible role of topological structure in the representation of large-scale navigable spaces. Third, there is work suggesting that cognitive maps might have a graph-like structure (Chrastil & Warren, 2014; Warren et al., 2017) as opposed to a coordinate-based, Euclidean structure (Gallistel, 1990; O’Keefe & Nadel, 1978; see also Yousif & Keil, 2021; Yousif, 2022). We view these more recent suggestions as consistent with earlier empirical work showing that angles in cognitive maps (revealed through drawing) are consistently biased (Byrne, 1979; Moar & Bower, 1983).

A topological framework can also help make sense of other distant findings. Take “shape skeletons”, for example. According to a now-orthodox view, objects are represented largely with respect to their internal skeletal structure (Ayzenberg & Lourenco, 2019; Firestone & Scholl, 2014; Lowet et al., 2018). While shape skeletons help to explain the underlying format of an object representation, they leave some questions unanswered. Namely: What parts of the shape skeleton do we represent (see Ayzenberg et al., 2019)? For certain complex objects, representing every detailed ridge via an additional skeletal branch would be computationally inefficient. So how does the visual system decide which branches contain the most meaningful information? Topology may be a critical part of the answer. That is: Certain segments may be more critical to the overall topological structure than others — and it may be those that are more likely to be preserved in the object representation. Indeed, this is one way of understanding our finding that T-junctions are preferentially preserved over L-junctions in the present tasks.

Chen (2005) argues that topological structure serves as a foundational structure for representation, and therefore “topological perception … is prior to the perception of other geometrical properties” (p. 3). This hierarchy is based upon the invariance of different systems to transformations, with topology being the most resilient, and consequently being encoded at a more “primitive” level than Euclidean features. This view also accords with a classic (yet unheralded) view in developmental psychology that children’s spatial representations are fundamentally topological in nature (Piaget & Inhelder, 1948). Indeed, a topological perspective can help to unify disparate ideas in vision science (Chen, 2005), developmental psychology (Piaget & Inhelder, 1948), and spatial navigation (Chrastil & Warren, 2014; Warren et al., 2017). The present work may provide a bridge between all of these research programs.

### “Core” knowledge of topology?

Topological knowledge interacts with other domains of knowledge in interesting ways. Topology is relevant not only for how we understand places and forms (as studied here), but also objects (Chien et al., 2012; Kibbe & Leslie, 2016; Kenderla et al., 2023) and number (Franconeri et al., 2009; He et al., 2015). In other words: Topology constitutes a kind of knowledge so foundational that it is critical to several domains of “core knowledge” (Spelke & Kinzler, 2007; Spelke, 2022). Indeed, in all of the cases cited above, topology seems to have an influence early in development (see, e.g., Clarke et al., 2025; Kibbe & Leslie, 2016; Yousif et al., 2025).

The fact that similar topological biases are observed in drawing tasks of large-scale spaces (Byrne, 1979; Moar & Bower, 1983) as well as small spaces (as shown here) may suggest a heretofore underappreciated commonality in the representation of forms and places — the fact that both may depend in part on topological primitives. If true, this would suggest that the accepted division between these two supposedly distinct domains may be more fragile than previously known, raising a host of new cognitive and developmental questions (see Garro-Rivero et al., 2026).

While we found ample evidence of topological biases in children, we did not find any definitive evidence that children were *more* dependent on topological structure than adults, perhaps inconsistent with the suggestion of Piaget and Inhelder (1948). Indeed, these findings are more consistent with the recent suggestion that topological intuitions are highly stable throughout development (Yousif et al., 2025).

### Conclusion

Every memory we make is full of errors. But if you look closely, these errors tell a story. Here, errors made in these drawing tasks demonstrate that people do not encode structures iconically, but with respect to some structured primitives. The present work moves us a step closer to understanding what those primitives are and, thus, a step closer to understanding the building blocks that support humans’ extraordinary ability to represent and manipulate the spatial world around them.

## Methods

### General

Sample sizes, dependent variables, and statistical tests for this experiment and all other experiments reported in this paper were determined and pre-registered prior to data collection (except Experiment S1, which was conducted in response to reviewer feedback). Preregistrations, stimuli, raw drawings, and data can be accessed on the following OSF page: https://osf.io/c26zj/overview

### Experiment 1

#### Participants

Fifty participants were recruited online via Prolific. All participants were eighteen years or older. They were residents of the United States who were fluent in English. Participants were excluded from the study if more than 30% of their data (i.e., more than three of the ten drawings) were unusable. This means the drawings either did not resemble the original stimulus at all, had topology/angles that were immeasurable, or they were determined by multiple members of the analysis team to be nonsensical and due to participant error. Two participants were excluded and replaced.

#### Stimuli

We created ten distinct stimuli. Five stimuli had holes and five stimuli did not have holes. The topologies of the stimuli were further varied by differing the number of T-junctions and L-junctions, with the simplest stimulus having only one L-junction and one T-junction, and the most complex stimulus having three T-junctions, three L-junctions, and one hole. The number of T-junctions and L-junctions were equated for all stimuli to allow us to more straightforwardly ask whether people are more likely to preserve T-junctions, L-junctions, or neither. In terms of angles, none of the stimuli included a 90° angle, as it could be used as a key reference when drawing and bias our data. The stimuli also did not include angles smaller than 20° or greater than 160° to prevent excessively sharp angles in the resulting drawings which could be perceived as single lines. Otherwise, we imposed no constraints on how T-junctions vs. L-junctions were incorporated (partially because these stimuli were based on earlier stimuli that were designed long before we imagined their use in this way; see Yousif & Brannon, 2024, 2025).

#### Design & Procedure

Participants were introduced to a memory task, in which they would see some objects and would need to draw them from memory. For each trial, an object would be shown to the participant for five seconds. The participant was instructed to memorize the object as well as they could. After the image disappeared, there was a five-second delay before the participant would be allowed to draw the object. Participants could draw by simply clicking and dragging the mouse. There was an “Undo” button to remove the latest stroke and a “Clear” button to clear the canvas entirely. At default zoom, the initial image and the subsequent canvas were each 600 pixels by 600 pixels in size. There was no time limit imposed during the drawing period. This process was repeated nine more times for a total of ten trials. Stimuli were randomized fully for each participant. Participants completed two representative practice trials before beginning the task. The practice trials were conducted on two separate stimuli, and the data/drawings from those trials were not recorded.

#### Analysis

Prior to detailed analysis, the drawings were first vetted superficially for any obvious participant errors, such as blank drawings due to technical errors or nonsensical drawings. Next, a team of six people manually analyzed each drawing. The drawings were separated by stimulus and assigned to different team members. Every drawing was coded independently by two different team members. First, every drawing was evaluated according to the exclusion criteria. In addition to checking once more for blank or nonsensical drawings, team members also evaluated whether topological features and angles could be measured. If a drawing had distinguishable topological features but lacked corresponding, measurable angles, the drawing was used in the analysis of only topological features. If a drawing lacked both, it was entirely excluded. To measure topological features (e.g., T-junctions), team members simply counted the L-junctions and T-junctions present in a drawing. Angle measurements were taken using an online measurement tool. Team members measured angles in the drawings which corresponded to those labelled in the stimuli. An absent angle was not scored. Angles were flagged for team review if there was uncertainty about whether a given junction was present (i.e., because it was hardly visible, or in a different location). After the initial coding phase, the two people who coded a single stimulus discussed and resolved any discrepancies, often with input from other team members. Any difference in the number of L-junctions and T-junctions in a drawing, as well as a difference of more than 10° in angle measurements, were automatically flagged for discussion. In the absence of a dispute, the final angle value used in the analysis was the average of the two coders’ values; if there was a dispute, both coders would re-evaluate and re-measure their original measurements, and one or both would adjust their value.

Based on the final angle values, we calculated bias towards (or away from) 90°. To do so, we simply calculated the initial distance away from 90°. An angle of 100° would have a difference of 10°, for example. Then we calculated the drawn distance away from 90°. An angle of 70° would have a difference of 20°. In a case like this, we would say that the drawn angle was 10° *further* than the original angle; thus this would constitute a bias away from 90°. Note that we count such cases as biases away from 90° despite the fact that the drawn angle actually went in the direction of 90°.

### Experiment 2

Everything about the design of this experiment was identical to Experiment 1, except as noted. 50 new participants completed the study. 7 additional participants were excluded. First, our stimuli for this experiment had only T-junctions and L-junctions, with no holes. Furthermore, the stimuli were more complex than the previous stimuli in that the range of topological features was from 2 L-junctions and T-junctions to 4 L-junctions and 4 T-junctions. We also designed six out of ten of the stimuli to have lines with lengths of a specific ratio. In these six critical stimuli, three of the lines had a 4:2:1 ratio, in which one line was twice the length of the shortest line, and another was four times the length of the shortest line. As with the angular analyses in Experiment 1, we planned to analyze whether the line lengths were distorted towards a central value (such that the drawn ratios might be closer to 3:2:1.5).

The analysis procedure was identical to the first experiment, except that (a) the line length analyses were restricted to the six critical stimuli with the 4:2:1 length ratios, and (b) in analyzing line lengths, we took the geometric mean of the ratio between the longest line and the middle line, and the middle line and the shortest line.

### Experiment 3

Everything about the design of this experiment was identical to Experiment 1, except as noted. 100 participants completed the task (see details below). Half of the stimuli were borrowed from Experiment 1 and half were borrowed from Experiment 2. Some of the stimuli were tweaked slightly to allow for easier visualization and analysis. Rather than merely having one large group of participants remember and then draw each of the ten drawings, we recruited ten participants at a time, in stages. In the first stage, ten participants completed the task, each remembering and then drawing the items in a random order. In the second stage, ten new participants were recruited. They would receive as input the drawings from the previous stage. They were additionally told that all lines were intended to be straight lines (to avoid misinterpretations of slight deviations). The chains were all intermixed so that each participant at each subsequent stage received one drawing from each participant of the previous stage (and, thus, never received more than one drawing from any one prior participant). For instance, the first participant in the second stage would see the first item from the first participant of the first stage, the second item from the second participant, and so on. The second participant in the second stage would see the second item from the first participant of the first stage, the third item from the second participant, and so on. Each subsequent stage proceeded with the same intermixed design, until a total of ten stages were completed. Given that ten participants created ten drawings in each of ten distinct stages, we were left with a final total of exactly 100 participants and 1000 analyzable drawings.

Throughout the data collection process, we removed and replaced any participant who provided nonsensical drawings (i.e., of the sort that would have been captured by the exclusion criteria of the previous experiments). Given the open-ended nature of the task, small disruptions at earlier stages of a chain would lead to uninterpretable data by the end. Thus, we had to be careful to avoid such disruptions, while also not biasing the chains to proceed in any particular direction. If we excluded any drawing from a participant, we excluded all of them and fully replaced that participant in the chain. In total, there were 97 exclusions of this sort. This means that of the approximately 2000 drawings we received in total, fewer than 5% were unusable. However, because we excluded any participant with a single failed drawing, this means a relatively high proportion of participants were excluded. Excluded drawings are shown in the materials on our OSF page.

To simplify our analyses, we focused our analyses primarily on the final 100 drawings (rather than on the 900 intermediate drawings).

### Experiment 4

The design of this experiment was identical to Experiment 1, except as noted. Data were collected from children at the Museum of Life and Science in Durham, North Carolina. We planned to collect data from 100 children (5-, 6-, 7-, and 8-year-olds, 25 from each age group). Unexpectedly, however, we found that five-year-old children were greatly underrepresented at the museum, such that, many months after data collection for the rest of the sample concluded, we were not close to completing the youngest group. In the meantime, we continued to collect data from the older groups. Rather than throw out these unexpected data, we retained them, in order to maximize the number of drawings available for analysis. Thus, our final sample consisted of 126 children (15 5-year-olds; 33 6-year-olds; 45 7-year-olds; and 33 8-year-olds; 45 children were excluded because an insufficient number of their drawings were usable). None of our conclusions depend on the inclusion of the additional children; we have included these data as part of an effort to maximize the number of drawings we are releasing publicly for purposes of future analysis.

The task itself was identical except for a few superficial changes. First, children were introduced to Leo the Lion and asked to help with his mission by completing the task. They were told that he was “exploring the galaxy” and had encountered some “mysterious shapes” that they needed to “help him remember”. Completing trials resulted in children being rewarded with a gem (a .gif of a flashing ruby, sapphire, or emerald). Second, children made their drawings using a stylus on a tablet computer rather than a mouse. Third, children were allowed to study the drawings for as long as they liked before the drawing phase. This was to account for external distractions that could prevent children from encoding the items within a fixed window. Fourth, an experimenter actively oversaw the children as they were drawing, allowing them to occasionally flag drawings to be discarded prior to data analysis. This was done to account for cases where the experimenter observed a clear distraction within a trial or a drawing that was going to be obviously unusable. Finally, the exclusion criterion for children was less rigorous. Data from an individual participant would be excluded only if more than 60% of drawings (i.e., seven or more out of ten) were deemed unusable.

